# HUH-Enabled Programmable Antibody-DNA Conjugates for Profiling Receptor-Specific Cellular Mechanics

**DOI:** 10.64898/2026.08.31.746762

**Authors:** Matthew R. Pawlak, Lidia K. Limon, Andrew M. Baldys, Matthew C. McMahon, Andrew C. D. Lemmex, Joseph M. Muretta, Frank M. Cichocki, Wendy R. Gordon

## Abstract

Cell-surface receptors integrate biochemical identity with mechanical information, yet methods for measuring receptor-specific force transmission across cell populations remain limited by the difficulty of coupling diverse recognition reagents to nucleic-acid tension probes. Here, we establish HUH endonuclease chemistry as a modular interface between antibodies and DNA-based mechanosensors. We develop two complementary strategies: genetically encoded HUH- antibody fusions that generate site-defined antibody-oligonucleotide conjugates in a single reaction, and a photocrosslinkable Protein G-HUH adaptor that enables covalent attachment of existing antibodies to DNA probes. Both approaches preserve antibody recognition while providing a programmable nucleic-acid handle for Rupture and Deliver Tension Gauge Tethers (RAD-TGTs). Using antibodies against beta1 integrin and HER2, we resolve receptor-specific mechanical phenotypes across cancer cell lines. Combining orthogonal features of beta1-integrin engagement and HER2 mechanical heterogeneity provides greater discrimination among cell types than either measurement alone, demonstrating that multidimensional mechanical phenotypes contain information not captured by individual force measurements. We further extend the platform to DNA:PNA tension probes to mitigate extracellular nuclease degradation and use antibody-functionalized RAD-TGTs to quantify force-dependent receptor engagement and pharmacological responses in immune cells. Together, these studies establish a modular antibody-to-nucleic-acid interface that expands DNA-based tension sensing beyond a restricted set of ligands and enables quantitative, multidimensional profiling of receptor-specific mechanical behavior across heterogeneous cell populations.

## Introduction

Antibody-oligonucleotide conjugates (AOCs) combine the molecular recognition of antibodies with the programmability of nucleic acids and are increasingly used for imaging, detection, targeted delivery, and molecular assembly.^1,2^ Their broader utility, however, depends on methods that connect antibodies to oligonucleotides with defined stoichiometry and geometry while preserving antigen recognition. This requirement becomes particularly stringent when the oligonucleotide is not simply a label, but a functional component of a molecular sensor.

DNA-based molecular tension probes provide one such application. By placing a receptor ligand on a DNA duplex with a defined rupture geometry, these probes report whether receptor-mediated forces exceed a specified molecular threshold. Tension gauge tethers and related sensors have revealed forces associated with integrin, Notch, and other mechanosensitive receptors,^3,4^ while Rupture and Deliver Tension Gauge Tethers (RAD-TGTs) convert probe rupture into delivery of a fluorescent oligonucleotide that can be quantified by flow cytometry.^5,6^ This enables force-threshold measurements across large cell populations. A major limitation, however, is the need to attach each receptor-targeting ligand to the nucleic-acid probe in a mechanically robust and experimentally tractable manner.

HUH endonucleases offer an attractive solution because they form sequence-directed covalent bonds with short single-stranded DNA substrates.^7,8^ HUH-tags can therefore provide a genetically encodable and chemically simple interface between proteins and nucleic acids. Here, we use HUH chemistry to develop two complementary routes for incorporating antibodies into DNA-based mechanosensors. In the first, a HUH domain is genetically fused to a recombinant antibody to provide a defined oligonucleotide attachment site. In the second, a photocrosslinkable Protein GHUH adaptor enables existing IgGs to be coupled covalently to DNA without engineering the antibody itself. During preparation of this work, Merkx and coworkers independently reported photocrosslinkable Protein G-HUH fusion proteins for generation of antibody-DNA conjugates for proximity extension assays.^9^ The work presented here extends this chemistry to mechanically loaded receptor-ligand interfaces and combines it with a genetically encoded HUH-IgG strategy for receptor-specific mechanophenotyping.

We apply these AOCs to DNA tension probes to ask whether multiple receptor-specific mechanical measurements can define cell states more effectively than individual measurements alone. Using beta1 integrin and HER2 as complementary receptor systems, we profile low- and high-threshold probe engagement and population heterogeneity across cancer cell lines and combine orthogonal features into a two-dimensional mechanical phenotype space. We further use the platform to quantify the concentration-dependent effects of LFA-1 modulators in primary human immune cells. Finally, because extracellular and membrane-associated nucleases can compromise DNA-based probes, we evaluate DNA:PNA hybrids as a nuclease-resistant alternative.^10,11^ Together, these studies establish HUH chemistry as a modular antibody-to-nucleic-acid interface and demonstrate how receptor-specific tension measurements can be combined into multidimensional mechanophenotypes as well as measure mechanokinetics.

## Experimental Methods

### Recombinant antibody production and HUH-mediated conjugation

Recombinant antibodies recognizing β1 integrin (K20) or HER2 (pertuzumab) were engineered with a C-terminal Wheat Dwarf Virus (WDV) HUH endonuclease tag on the antibody heavy chain. Heavy- and light-chain plasmids were transiently co-transfected into Expi293F cells, and secreted antibodies were purified from conditioned medium by Protein A affinity chromatography. HUH-mediated DNA conjugation was performed by incubating HUH-tagged antibodies with single-stranded DNA substrates containing the WDV recognition sequence in 50 mM HEPES (pH 8.0), 50 mM NaCl, and 1 mM MnCl_2_ for 30 min at 37 °C. Conjugation efficiency was assessed by SDS-PAGE. Antigen binding of HUH-tagged antibodies was confirmed by biolayer interferometry using recombinant β1 integrin or HER2 ectodomains. Detailed cloning, expression, purification, conjugation, and binding protocols are provided in the Supporting Information.

### Photocrosslinkable Protein G–HUH adaptor

To enable HUH-mediated modification of antibodies without genetic engineering, the Fc-binding domain of Protein G was fused to WDV HUH and engineered to incorporate the photocrosslinkable amino acid *p*-benzoyl-L-phenylalanine at Ala24. The resulting clPG-HUH adaptor was expressed in *E. coli*, purified, incubated with Fc-containing antibodies, and irradiated at 365 nm to generate covalent antibody–HUH conjugates. Formation of the covalent complex and retention of HUH conjugation activity were evaluated by SDS-PAGE.

### Preparation of receptor-specific RAD-TGTs

DNA tension-gauge-tether (TGT) duplexes corresponding to low- and high-force rupture thresholds were prepared by annealing complementary oligonucleotides and subsequently coupled to HUH-tagged proteins through the sequence-directed HUH reaction. For receptor-specific mechanophenotyping, K20-HUH and pertuzumab-HUH were coupled to TGTs to generate β1-integrin- and HER2-targeting probes, respectively. Glass-bottom plates were sequentially functionalized with biotinylated BSA, neutravidin, and biotinylated RAD-TGT probes. Cells were plated onto functionalized surfaces, where receptor engagement and cell-generated forces exceeding the rupture threshold separated the DNA duplex and transferred a fluorescent oligonucleotide to the cell.

### Flow-cytometric mechanophenotyping

Following incubation on RAD-TGT surfaces, cells were recovered and analyzed by flow cytometry with the gating strategy shown in Figure S8. Receptor-specific mechanical phenotypes were quantified from the fraction of RAD-TGT-positive cells and the fluorescence intensity and heterogeneity of the delivered oligonucleotide signal. For multidimensional analysis of U251, SKBR3, BT474, and SKOV3 cells, β1-integrin low-threshold engagement and HER2 high-threshold signal heterogeneity were combined to generate a twodimensional mechanical phenotype. Pairwise separation between cell lines was calculated after standardization of individual features, with two-dimensional separation calculated from the Euclidean distance between the two coordinates. Statistical significance of the improvement in separation obtained by combining the two receptor measurements was evaluated by permutation testing and bootstrap resampling as described in the Supporting Information.

### LFA-1 mechanical pharmacology

Primary human NK cells were treated with LFA-1-modulating antibodies and subsequently plated on ICAM-1-functionalized RAD-TGT surfaces. Delivered fluorescent oligonucleotide was quantified by flow cytometry as a direct measure of changes in receptor-mediated force transmission. Concentration-response curves were fit by nonlinear regression, and concentrations producing half-maximal increases or decreases in the mechanical response were designated mechanical EC_50_ (mEC_50_) or IC_50_ (mIC_50_), respectively.

### Surface nuclease and DNA:PNA RAD-TGT assays

Surface nuclease activity was measured using fluorophore/quencher DNA probes immobilized on glass surfaces, with probe cleavage visualized by fluorescence microscopy and quantified using CellProfiler. DNA:PNA tension probes were prepared by annealing complementary DNA and PNA strands and evaluated using HUH-echistatin-functionalized RAD-TGTs. Effects of ROCK inhibition with Y-27632 on DNA:DNA and DNA:PNA probe rupture were quantified by flow cytometry. Complete probe sequences, experimental conditions, imaging procedures, and statistical analyses are provided in the Supporting Information.

## Results and Discussion

### HUH-tags provide complementary routes for coupling antibodies to DNA

To create a modular connection between antibodies and nucleic-acid probes, we developed two complementary HUH-tag-based strategies (Figure 1). HUH endonucleases recognize short singlestranded DNA sequences and form a covalent phosphotyrosine linkage to the cleaved DNA, enabling rapid, sequence-directed protein-DNA conjugation.^7,8^ In the first strategy, a Wheat Dwarf Virus HUH-tag was genetically fused to the C-terminus of the heavy chain of full-length IgGs targeting beta1 integrin or HER2 (Figure 1B, Figure S1). Incubation with cognate single-stranded DNA produced efficient covalent conjugation, as assessed by SDS-PAGE. Importantly, incorporation of the HUH domain and subsequent DNA conjugation preserved target recognition: HUH-tagged and corresponding commercial antibodies showed similar recognition patterns across cancer-cell lysates, and biolayer interferometry measurements showed comparable binding affinities (Figure 1C,D; Figure S2) for the recombinant and untagged reagents. These results support genetically encoded HUH fusion as a direct route to antibody-DNA conjugates with a defined oligonucleotide attachment site.

**Figure 1.**
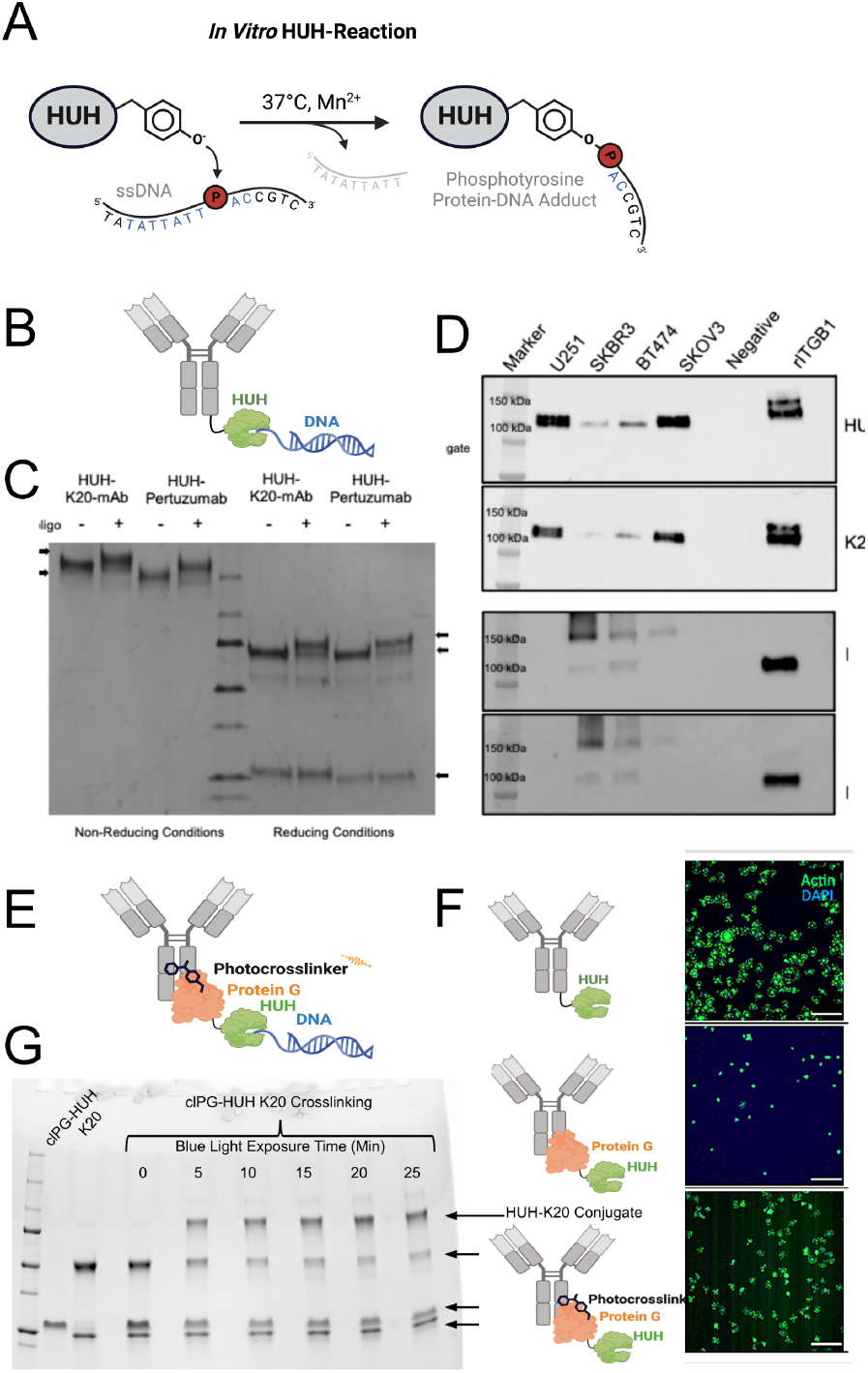
Complementary HUH-based strategies for coupling antibodies to oligonucleotides. (A) HUH endonucleases recognize cognate single-stranded DNA sequences and form covalent protein-DNA adducts. (B) Direct strategy in which a HUH-tag is genetically encoded at the C-terminus of an antibody heavy chain. (C) SDS-PAGE analysis of recombinant beta1-integrin- and HER2-targeting antibodies before and after reaction with cognate oligonucleotides. (D) Comparison of target recognition by HUH-tagged and commercial antibodies across cancer-cell lysates. (E) Adaptor strategy in which Protein G-HUH binds the Fc region of an existing antibody and is covalently trapped by Bpa-mediated photocrosslinking. (F) Representative cell-adhesion images comparing direct antibody-HUH probes, noncovalent Protein G-HUH attachment, and photocrosslinked Protein G-HUH attachment. (G) SDS-PAGE analysis of photocrosslinking and HUH-mediated oligonucleotide conjugation.

### A photocrosslinkable Protein G-HUH adaptor enables use of existing antibodies

Genetic fusion is attractive when recombinant antibody production is practical, but many experiments depend on existing commercial antibodies. We therefore developed an adaptor-based strategy in which a Protein G-HUH fusion connects an IgG to the DNA probe (Figure 1E). Protein G binds the Fc region of IgGs with high affinity and has previously been used to present Fc-containing ligands on molecular tension sensors.^12^ However, noncovalent Protein G-Fc attachment was insufficient under our assay conditions: cells showed substantially reduced adhesion to surfaces in which the antibody was connected to the probe only through the noncovalent Protein G-Fc interaction (Figure 1F), consistent with disruption of the complex under mechanically loaded conditions.

To stabilize this interface, we incorporated the photocrosslinkable amino acid Bpa at the Protein GFc binding interface, building on photocrosslinkable Protein G approaches for site-selective IgG conjugation.^13^ UV irradiation converted the Protein G-IgG interaction into a covalent complex while retaining HUH-mediated DNA conjugation (Figure 1G, Figure S4). Crosslinking increased with irradiation time, whereas prolonged UV exposure reduced HUH activity, defining an experimental window that balances the two reactions (Figure S5). During preparation of this work, Merkx and coworkers independently reported photocrosslinkable Protein G-HUH fusion proteins for production of homogeneous antibody-DNA conjugates used in proximity extension assays.^9^ Our implementation uses the same general Protein G-HUH concept in a mechanically loaded setting and, together with the direct HUH-IgG strategy, provides complementary engineered and adaptor-based routes for coupling antibodies to defined nucleic-acid probes.

### Antibody-functionalized RAD-TGTs generate receptor-specific mechanical phenotypes

We next incorporated the antibody-DNA conjugates into RAD-TGTs (Figure 2A). In RAD-TGT assays, receptor engagement and cytoskeletal force rupture a DNA duplex and release a fluorescent oligonucleotide that is subsequently detected by flow cytometry.^5^ Altering the duplex geometry produces low- and high-threshold probes (Figure S3), allowing receptor engagement to be examined under different mechanical stringencies. Flow cytometry additionally reports the distribution of signal across the population, providing information about both mean engagement and cell-to-cell heterogeneity.

**Figure 2.**
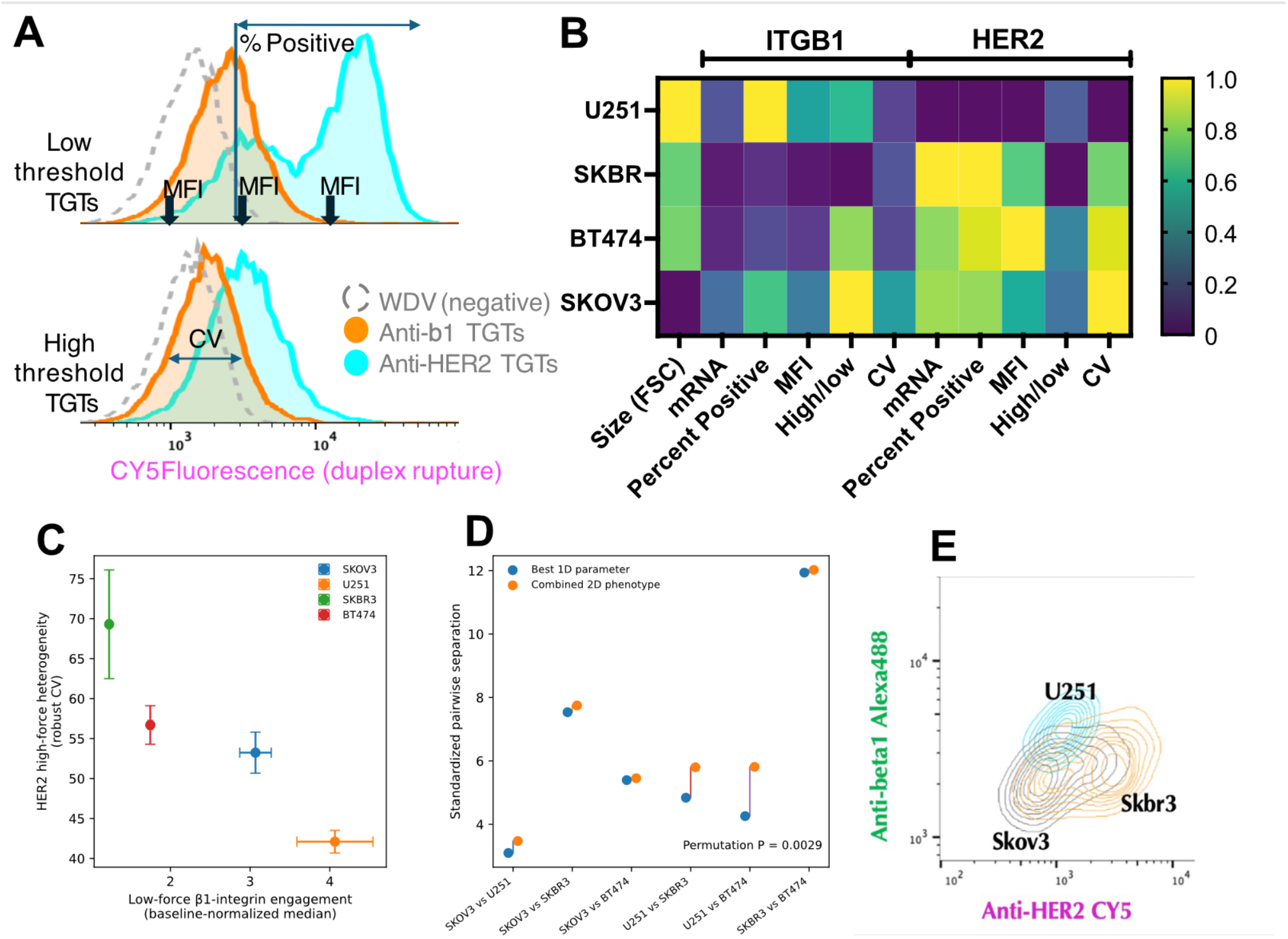
Antibody-functionalized RAD-TGTs enable multidimensional, receptor-specific mechanophenotyping of cancer cells. (A) Representative flow-cytometry distributions illustrating mechanical features extracted from cells plated on low- and high-threshold RAD-TGTs functionalized with antibodies against β1 integrin (ITGB1; orange) or HER2 (cyan). WDV-only TGTs lacking a targeting ligand serve as a negative control (gray dashed line). RAD-TGT rupture delivers a Cy5-labeled oligonucleotide to cells, enabling quantification of median fluorescence intensity (MFI), percentage of probe-positive cells, the ratio of highto low-threshold signal, and population heterogeneity measured by coefficient of variation (CV). **(B)** Heatmap summarizing receptor expression and RAD-TGT-derived mechanical features across U251, SKBR3, BT474, and SKOV3 cells for ITGB1 and HER2. Features include cell size (forward scatter, FSC), relative mRNA abundance, percentage of RAD-TGT-positive cells, MFI, high/low-threshold signal ratio, and CV. Values are scaled independently by feature from 0 to 1 to visualize relative differences among cell lines. **(C)** Two-dimensional mechanical phenotype space defined by normalized low-threshold β1-integrin engagement and HER2 high-threshold signal heterogeneity (CV). Points represent the mean of independent biological replicates; horizontal and vertical error bars indicate SEM. Combining measurements from two receptors resolves the four cancer cell lines into distinct regions of mechanical phenotype space. **(D)** Quantitative comparison of cell-line separation using the best-performing individual one-dimensional (1D) parameter (blue) versus the combined twodimensional (2D) phenotype shown in (C) (orange). Each pair of points represents one pairwise cell-line comparison. The combined 2D phenotype increased overall pairwise separation relative to the best individual parameter (**permutation P = 0.0029**). **(E)** Multiplexed receptor-specific RADTGT profiling. HER2- and β1-integrin-targeting RAD-TGTs labeled with spectrally distinct fluorophores were presented simultaneously, allowing receptor-specific mechanical engagement to be measured within the same cells by flow cytometry. Representative contour plots illustrate distinct two-dimensional β1-integrin/HER2 mechanical signatures for SKOV3, U251, and SKBR3 cells.

Using beta1-integrin- and HER2-targeting probes, we measured a panel of RAD-TGT features across BT474, SKBR3, SKOV3, and U251 cells. Low-threshold measurements captured substantial differences in receptor-dependent engagement across the lines, whereas high-threshold measurements imposed greater mechanical stringency. Importantly, no single feature fully resolved all four cell lines. This suggested that receptor-specific mechanical measurements might contain complementary information that is lost when each parameter is considered independently.

### Orthogonal receptor features define a two-dimensional mechanophenotype

We therefore compared pairs of candidate features and identified normalized beta1-integrin low-threshold engagement and HER2 high-threshold signal heterogeneity (CV) as complementary coordinates. Plotting these measurements together separated the four cancer cell lines in a twodimensional mechanical phenotype space (Figure 2C). Quantitative comparison of onedimensional and two-dimensional separation showed that the combined feature space discriminated the cell lines better than either selected feature alone. Thus, the value of the platform is not limited to measuring whether a receptor exceeds a particular force threshold; multiple receptor-specific measurements can be integrated to define higher-dimensional mechanical cell states.

HER2 provided a particularly informative heterogeneity dimension. HER2-high cell lines displayed broad, and in some cases visibly heterogeneous, high-threshold fluorescence distributions. We interpret this conservatively as evidence for distinct HER2 engagement states within the population. HER2 and beta1-integrin signaling are known to interact in cancer cells,^14,15^ raising the possibility that differences in receptor organization or coupling to adhesion machinery contribute to these states; however, the molecular basis of the observed heterogeneity remains to be established. The present data therefore identify HER2 high-threshold CV as a useful phenotypic feature rather than demonstrating a specific HER2-integrin mechanical coupling mechanism.

### Multiplexed RAD-TGT measurements can resolve heterogeneous populations

To further test whether receptor-specific mechanical information could be multiplexed, we generated RAD-TGTs in which HER2- and beta1-integrin-targeting probes carried distinct fluorophores. In mixed-cell experiments, dual-color flow cytometry resolved populations according to their receptor-specific probe signatures without requiring prior assignment of cell identity. These experiments support the broader use of antibody-functionalized RAD-TGTs for function-based phenotyping of heterogeneous cell populations and motivate expansion to larger receptor panels.

### RAD-TGTs quantify concentration-dependent effects of LFA-1 modulators

We next asked whether antibody-enabled RAD-TGTs could quantify pharmacological modulation of a mechanically regulated receptor (Figure 3). Because the low-threshold probe produced a larger dynamic RAD-TGT signal in primary NK cells than the high-threshhold probe (Figure S6), we used this probe for subsequent concentration dependent force measurements. Primary human NK cells were treated with antibodies that inhibit, activate, or report the activation state of LFA-1 before plating on ICAM-1-functionalized RAD-TGTs. TS1/18 produced a concentration-dependent reduction in RAD-TGT signal, whereas CBR LFA-1/2 produced a concentration-dependent increase over the activating range tested.^16,17^ These responses reproduced the known inhibitory and activating behavior of the reagents in a receptor-specific mechanical readout.

**Figure 3.**
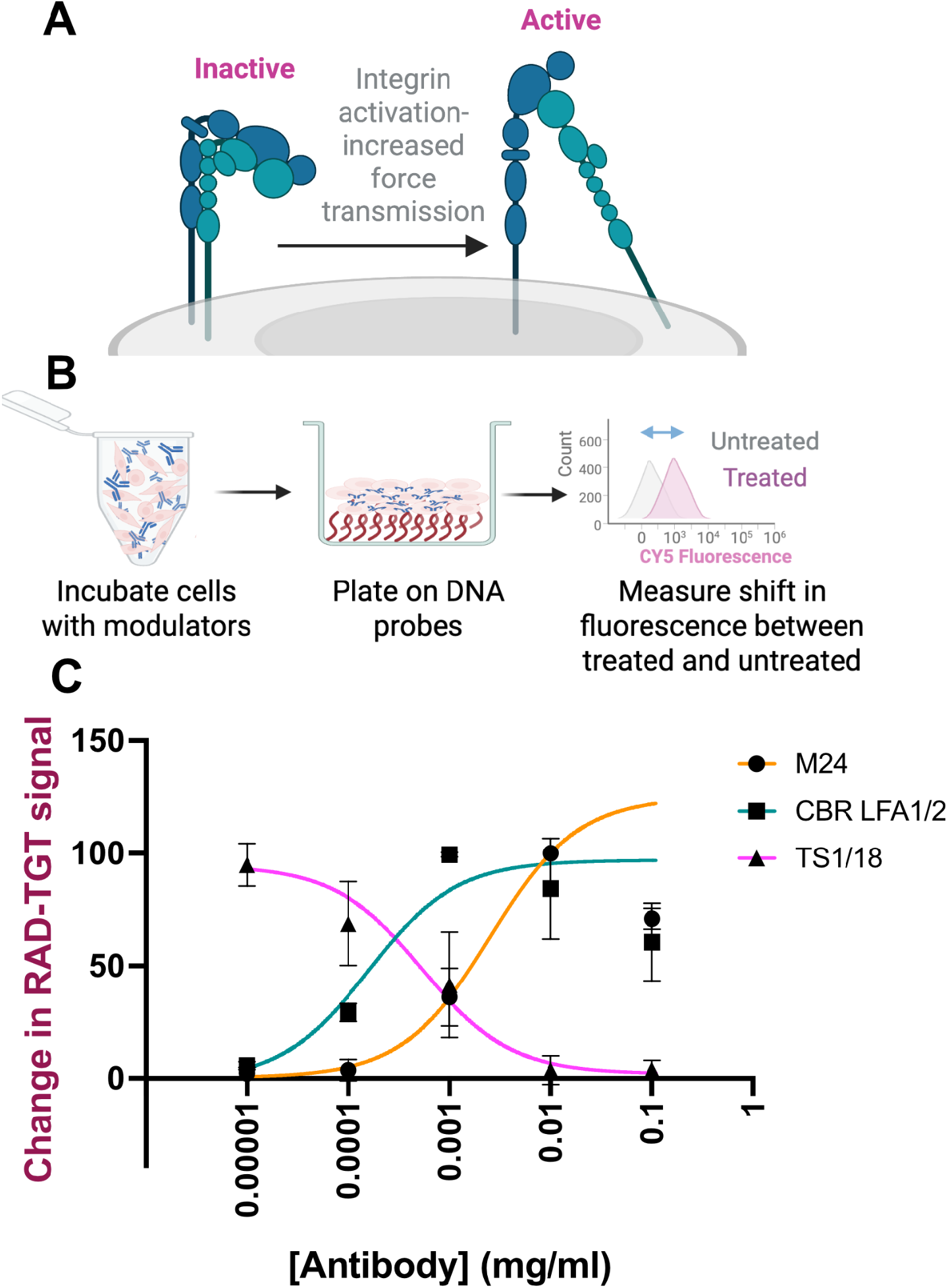
RAD-TGT measurement of concentration-dependent mechanical responses to LFA-1 modulators. (A) Schematic representation of LFA-1 conformational activation and mechanically productive ligand engagement. (B) Experimental workflow. Primary human NK cells were incubated with increasing concentrations of LFA-1 modulators and plated on ICAM-1-functionalized RAD-TGT surfaces; probe rupture and fluorescent oligonucleotide delivery were quantified by flow cytometry. (C) Concentration-response curves for TS1/18, CBR LFA-1/2, and M24. Values are reported as changes in RAD-TGT signal relative to the appropriate untreated condition. Apparent mEC50 or mIC50 values describe half-maximal changes in the mechanical RAD-TGT response and are not intended as conventional binding-affinity measurements.

To distinguish these measurements from conventional binding or signaling potency values, we describe concentrations producing half-maximal changes in RAD-TGT signal as mIC50 or mEC50 values. TS1/18 produced a robust inhibitory mechanical response with an apparent mIC50 in the low-nanomolar range. CBR LFA-1/2 produced an activating response at still lower concentrations. At higher concentrations, however, the RAD-TGT response decreased for some activating treatments (Figure 3C and Figure S7A), producing a bell-shaped concentration-response profile. Such behavior could arise from several mechanisms, including steric interference with LFA-1-ICAM-1 engagement, altered antibody-mediated receptor crosslinking at high occupancy, or stabilization of receptor states that are activated but less effectively coupled to ligand engagement and force transmission. Thus, maximal receptor occupancy need not correspond to maximal mechanically productive engagement. These profiles illustrate an advantage of the assay: RAD-TGTs report the mechanical consequence of receptor modulation rather than antibody binding or receptor activation alone. M24, which is commonly used as a reporter of the extended-open LFA-1 conformation, also altered the mechanical response. Its concentration-response profile was shifted relative to CBR LFA-1/2, consistent with weaker activation in this assay. Because M24 binds and stabilizes an activated/extended LFA-1 conformation, an effect on mechanically productive engagement is mechanistically plausible.^18,19^ We therefore interpret M24 as an activation-state reporter that can also perturb the conformational equilibrium under these assay conditions, rather than as a purely passive label. Similar experiments in U251 cells with RGD-binding integrin modulators produced ligand-dependent responses (Figure S7B), supporting the generalizability of RAD-TGT-based mechanical pharmacology across receptor and cell contexts.

### Nuclease-resistant probes extend the platform to challenging biological environments

DNA-based mechanosensors are vulnerable to extracellular and membrane-associated nucleases, which can reduce probe integrity independently of mechanical rupture.^11^ We therefore evaluated extracellular nuclease activity using a surface nuclease sensor (SNS, Figure S9) and DNA:PNA hybrid tension probes as a nuclease-resistant alternative.^20^ Using surface nuclease sensors, we observed contributions from both serum-associated and cell-associated nuclease activity, with particularly strong localized activity associated with C2C12 cells **(Figure S10)**. DNA:PNA probes produced lower absolute RAD-TGT signal than corresponding DNA duplexes, consistent with their greater duplex stability (Figure S11). In systems with elevated surface nuclease activity, DNA:PNA probes better preserved force-dependent signal and could recover mechanically induced changes that were obscured with DNA-only probes (Figure S12). These results establish nuclease resistance as an important design variable when deploying antibody-linked tension sensors across diverse cell types and provide a practical route for adapting the platform to high-nuclease environments.

## Conclusions

Together, these studies establish HUH chemistry as a modular interface between antibody recognition and nucleic-acid mechanosensors. Direct HUH-IgG fusions provide a genetically encoded attachment site, whereas photocrosslinkable Protein G-HUH adaptors enable existing antibodies and Fc-containing ligands to be incorporated into mechanically robust probes. Coupling these reagents to RAD-TGTs expands tension sensing from a restricted set of ligands toward receptor-specific, multiplexable measurements across cell populations. Most importantly, combining orthogonal measurements of receptor engagement and population heterogeneity creates multidimensional mechanophenotypes that contain discriminatory information not apparent from individual force measurements. The same framework can be used to quantify pharmacological perturbation and can be adapted with nuclease-resistant probe chemistries. We envision antibody-linked nucleic-acid mechanosensors as a general strategy for translating receptor identity, force threshold, and cellular heterogeneity into quantitative mechanical phenotypes.

