## Supplementary information for "HUH-Enabled Programmable Antibody-DNA Conjugates for Profiling Receptor-Specific Cellular Mechanics"

### Supplemental Figures

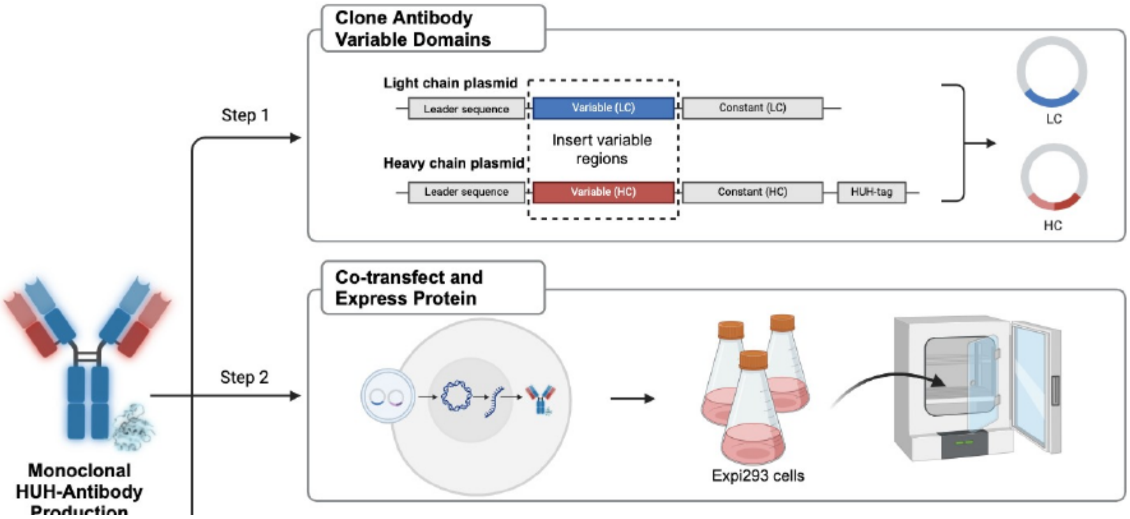


Supplementary Figure S1. Design of recombinant HUH-antibody constructs and initial expression strategy. HUH-tags were placed at the C terminus of either the antibody heavy or light chain to compare expression and DNA-conjugation activity. This source figure is retained as a candidate supplement because the combined manuscript focuses on the heavy-chain construct but does not show the initial construct-design comparison.

A

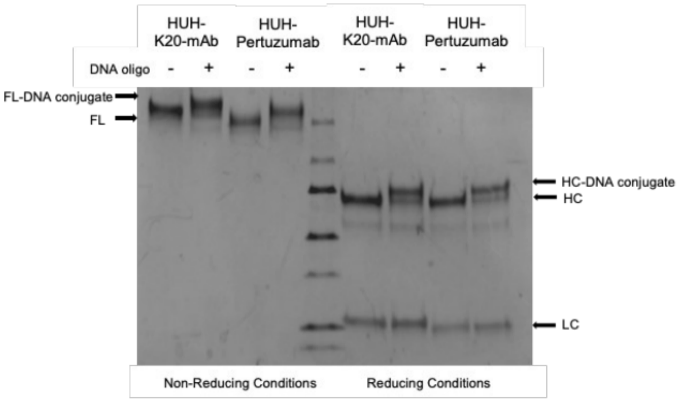


C

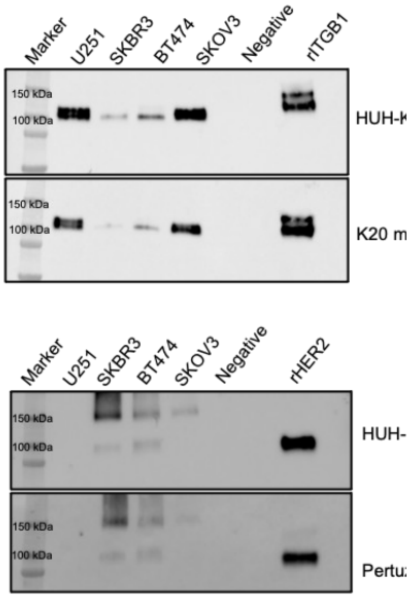


B

Binding affinity of anti-HER2 mAbs to its HER2 ligand and of anti-K20 mAbs to the integrin β1 ligand.

| Antibody | Affinity* (Kd [nM]) |
|---|---|
| Pertuzumab | 3.9 nM |
| HUH-Pertuzumab | 5 nM |
| K20 | 212 nM |
| HUH-K20 mAb | 134 nM |

* Affinity was measured by bio-layer interferometry using recombinant αVβ1 integrin heterodimer for K20 mAbs and ErbB2-ECD for Pertuzumab mAbs.

Supplementary Figure S2. Additional characterization of HUH-tagged K20 and pertuzumab antibodies. Source data include HUH-mediated oligonucleotide conjugation and biolayer-interferometry measurements comparing recombinant HUH-tagged antibodies with commercial counterparts. The BLI data are especially important because the combined manuscript states that binding affinities are similar but currently refers to a missing supplementary figure. Final panel lettering and numerical K_D values should be rebuilt from the original analysis files.

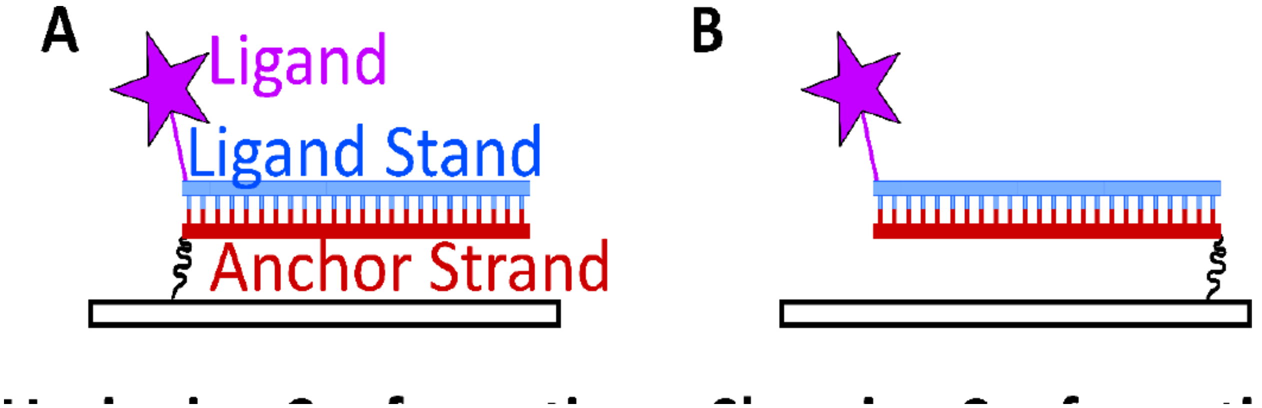


Supplementary Figure S3. Tension-gauge tether geometry. Positioning of ligand and surface anchor on proximal termini creates an unzipping geometry with a lower rupture threshold, whereas placement on distal termini creates a shearing geometry with a higher rupture threshold. Intermediate thresholds can be generated by changing anchor position.

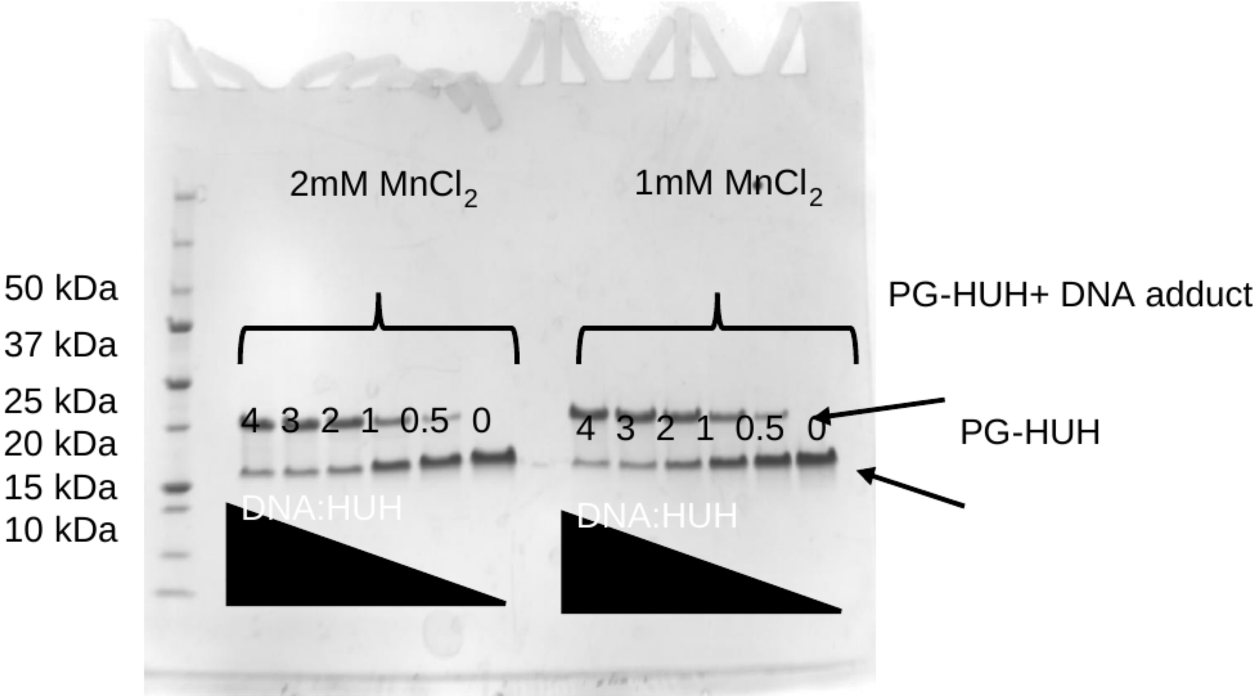


Supplementary Figure S4. Protein G-HUH retains HUH-mediated DNA conjugation activity. Source gel data show formation of a higher-molecular-weight PG-HUH-DNA adduct under HUH reaction conditions. This experiment supports use of PG-HUH as the DNA-coupling module in the indirect antibody-conjugation strategy.

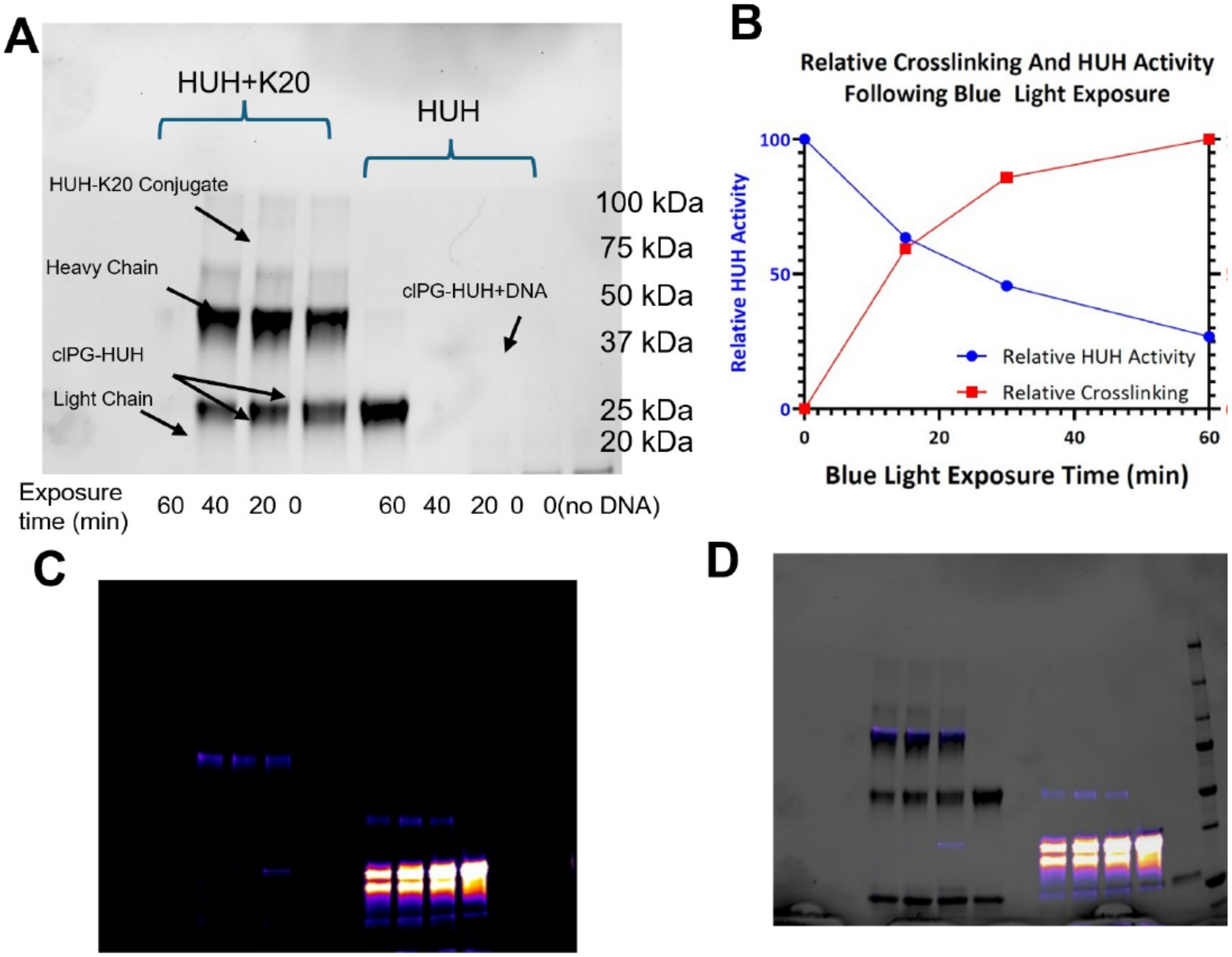


Supplementary Figure S5. Optimization of photocrosslinking of clPG-HUH to K20 while retaining HUH activity. The source manuscript tested increasing 365-nm illumination times and quantified the tradeoff between Fc crosslinking and subsequent oligonucleotide conjugation. Approximately 20 min illumination was selected as a practical compromise. The final SI should consolidate these source panels into a single clean multi-panel figure and replace image-derived quantification with plots exported from the original analysis files.

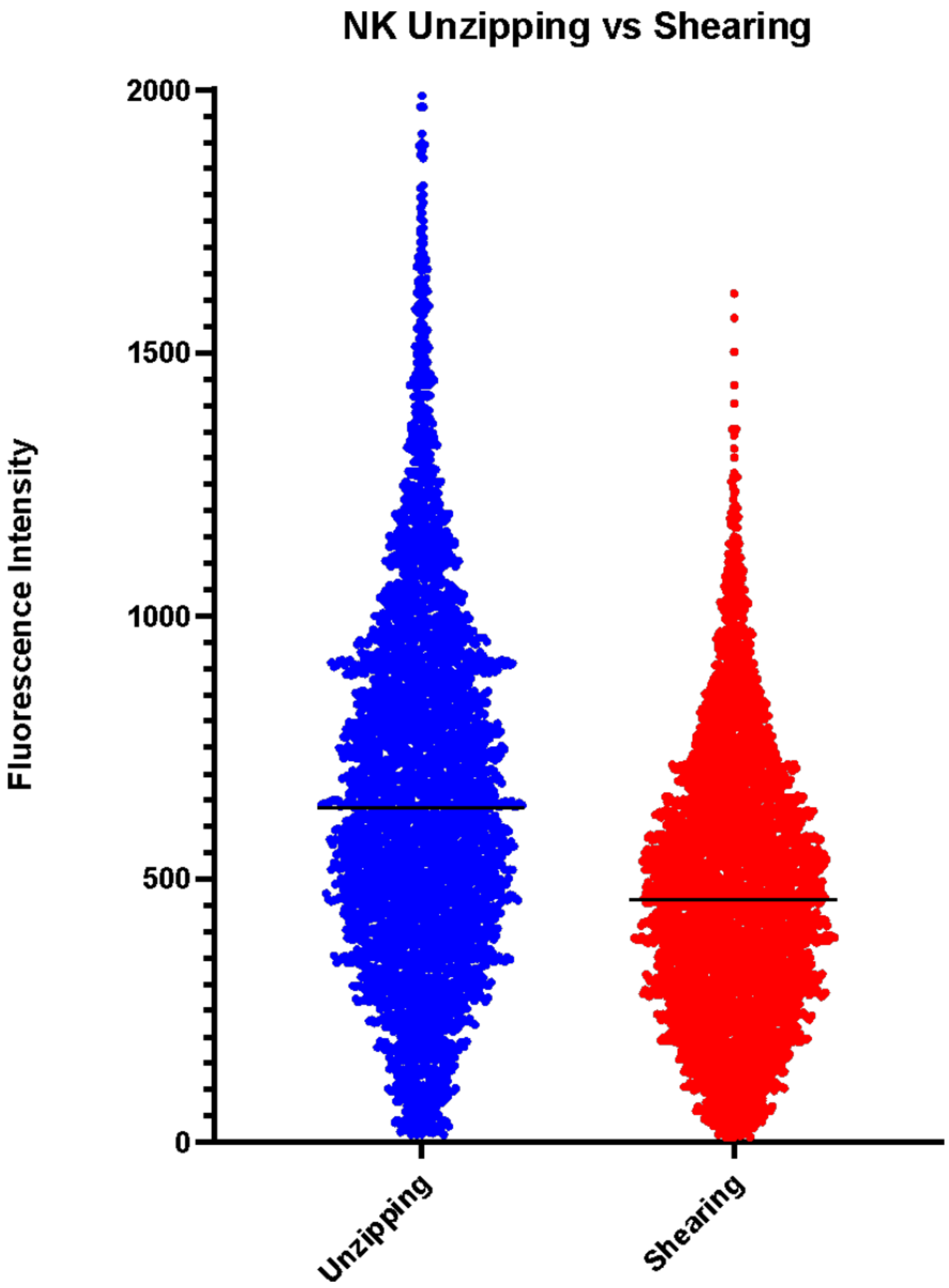


Supplementary Figure S6. Distribution of RAD-TGT fluorescence in primary NK cells plated on ICAM-1-presenting low-force (unzipping) or high-force (shearing) probes. This source experiment supports the choice of the low-force probe for the mechanokinetic titration because it provides greater signal in NK cells.

**A**

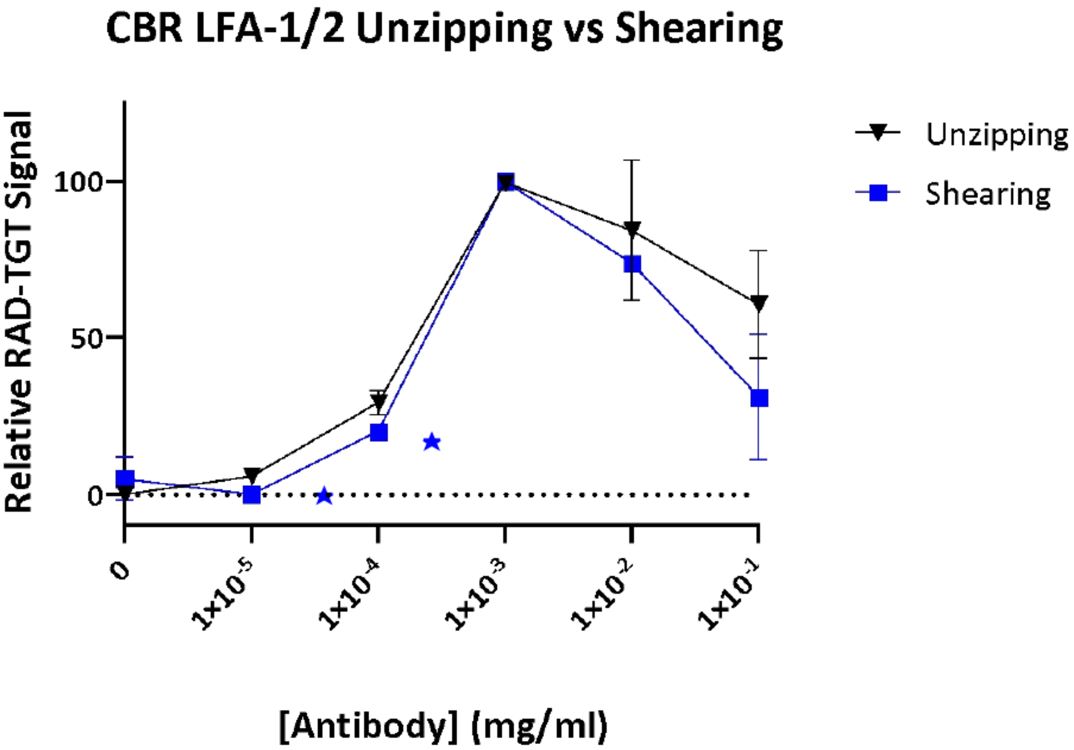


**B**

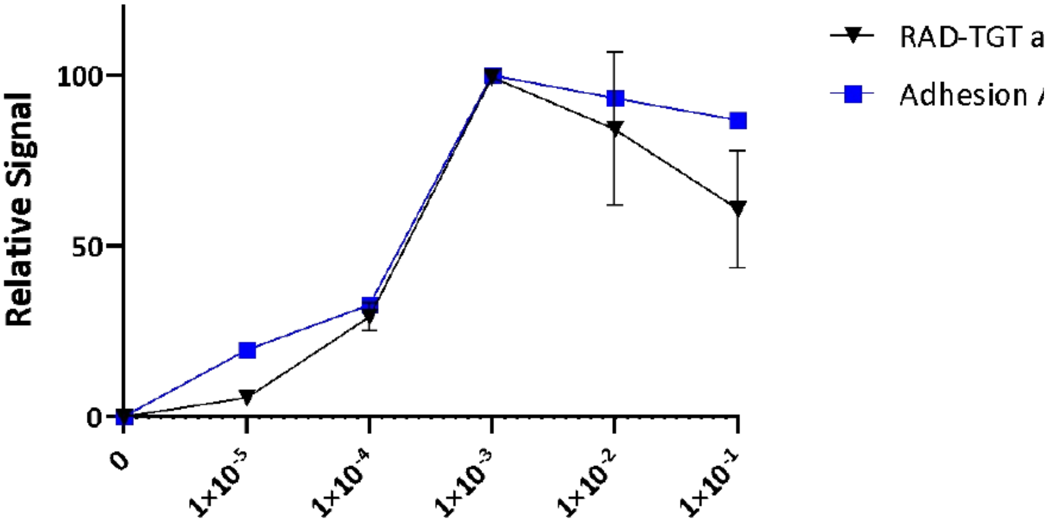


Supplementary Figure S7A. CBR LFA-1/2 dose response across complementary assay conditions. Source data compare the mechanokinetic response measured with low-force RAD-TGTs with measurements on high-force probes and with previously reported adhesion behavior. This provides context for the non-monotonic response observed at higher concentrations of activating antibody. Final use should be checked for permissions if any panel directly reproduces published literature data rather than replotted numerical values.

A

B

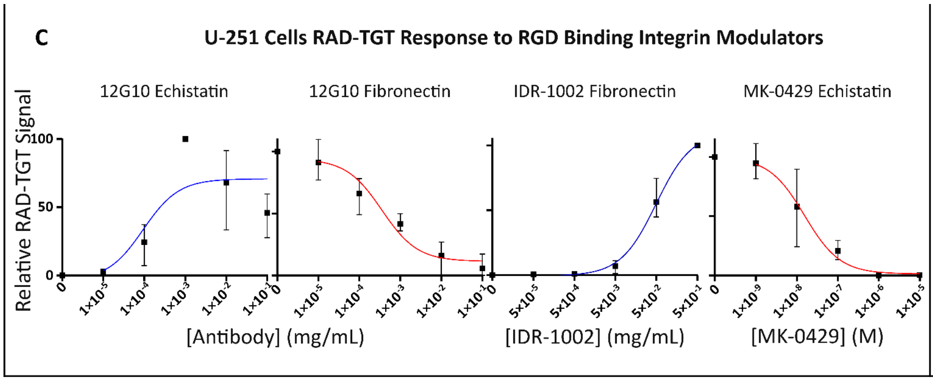


D

| Treatment | Ligand | EC50 [mg/mL] | |
|---|---|---|---|
| | | Value | SEM |
| IDR-1002 | Fibronectin | $4.9x10^{-2}$ | $9x10^{-3}$ |
| 12G10 | Echistatin | $1x10^{-4}$ | $1x10^{-4}$ |
| | | **IC50 [mg/mL]** | |
| 12G10 | Fibronectin | $4x10^{-4}$ | $2x10^{-4}$ |
| | | **IC50 [M]** | |
| MK-0429 | Echistatin | $1.54x10^{-8}$ | $6.09x10^{-9}$ |

**Supplementary Figure S7B : Mechanokinetic Assessment of Integrin Modulators:** A-B) Quantification of the effect of RGD binding integrin targeting drugs on U-251 cells using unzipping RAD-TGTs conjugated to either echistatin-WDV or fibronectin-WDV. Graphs of relative RAD-TGT signal of U-251 cells fitted with functional binding curves following titration of the drug indicated in title. The title also indicates ligands used for assessment. Data presented representing 3 independent experiments of >1000 cells. B) The EC50 or IC50 of the fitted curves in 5C and standard error and mean for each value.

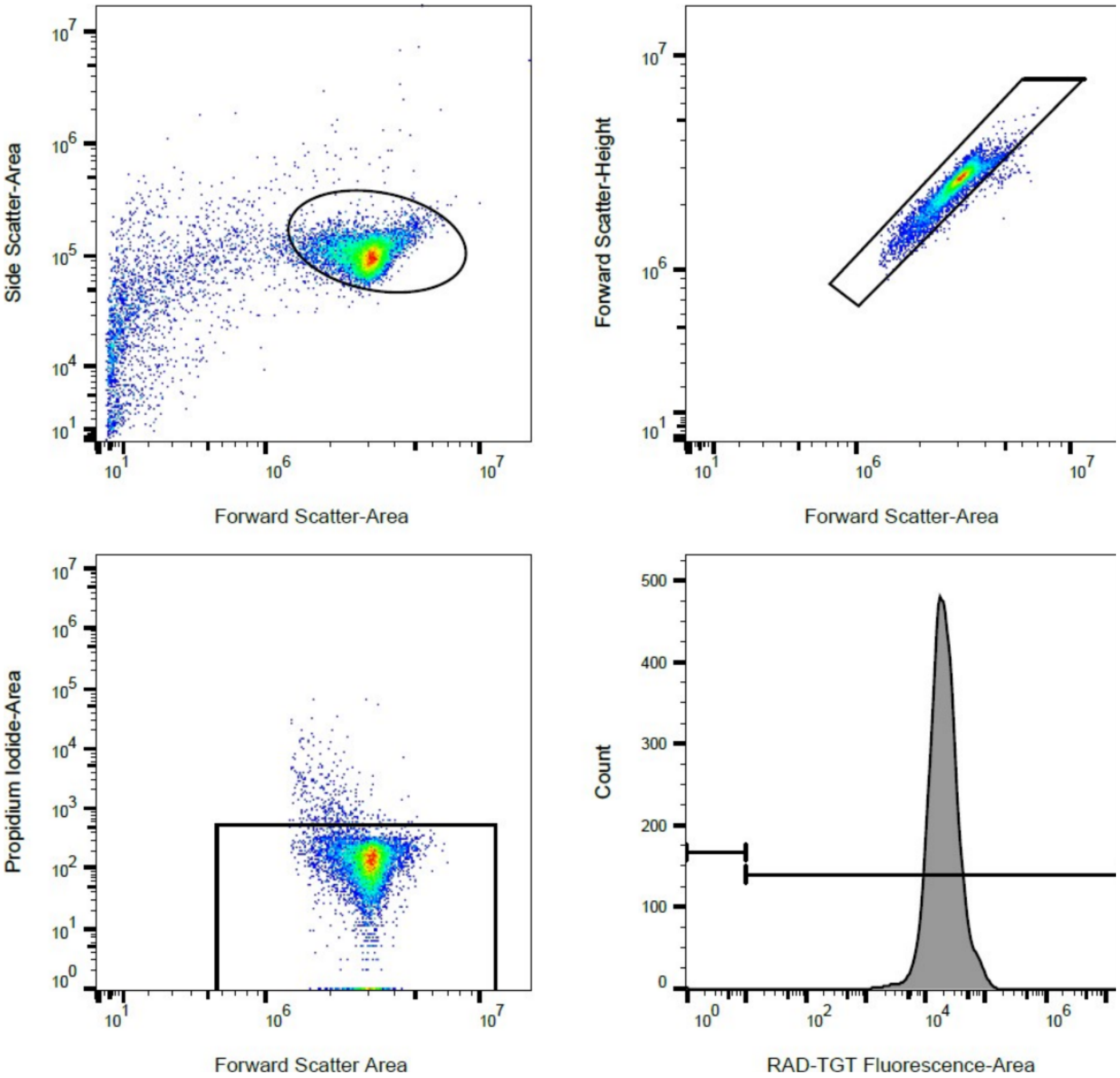


Supplementary Figure S8. Representative flow-cytometry gating strategy used for RAD-TGT experiments. Cells are selected by forward and side scatter, singlets are isolated using FSC-A versus FSC-H, viable cells are selected using propidium iodide, and the RAD-TGT fluorescence population is then quantified relative to the appropriate negative control.

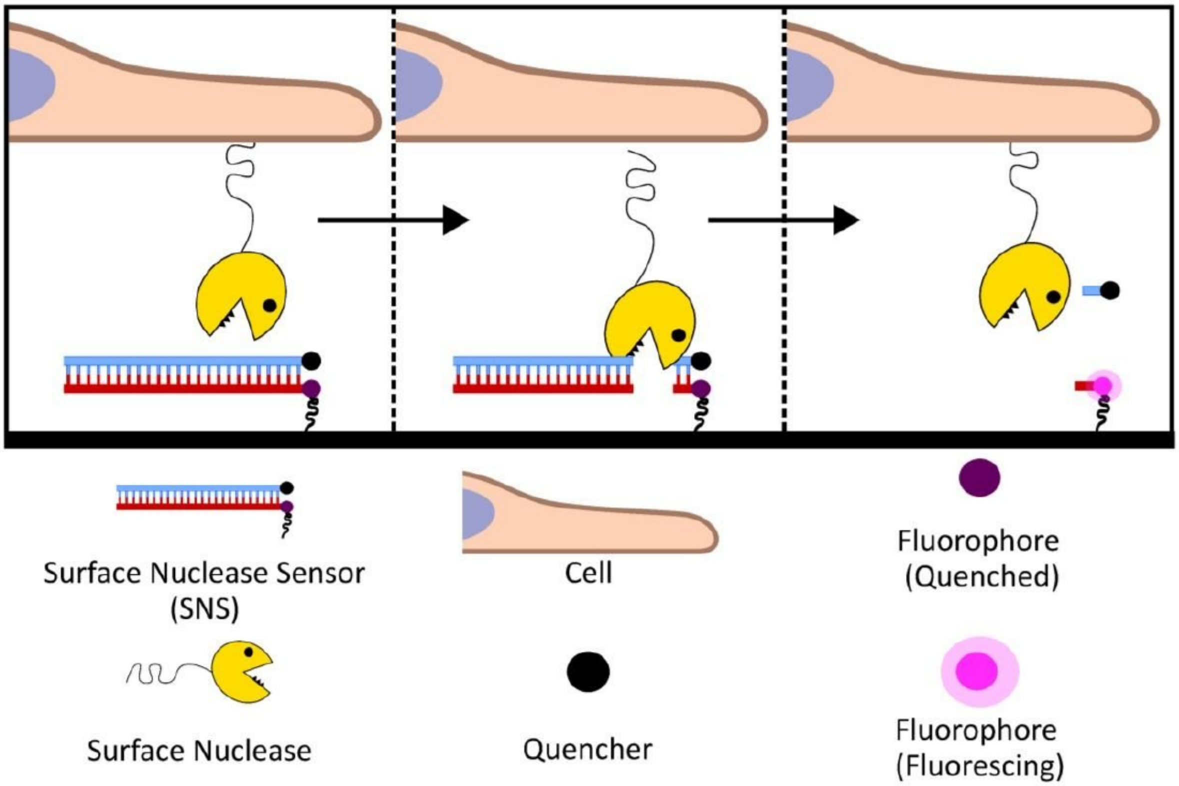


Supplementary Figure S9. Principle of the surface nuclease sensor (SNS). SNS probes are surface-immobilized DNA duplexes bearing a fluorophore-quencher pair but no receptor ligand. Nuclease cleavage destabilizes the duplex and separates fluorophore from quencher, generating a local fluorescent signal that reports extracellular nuclease activity.

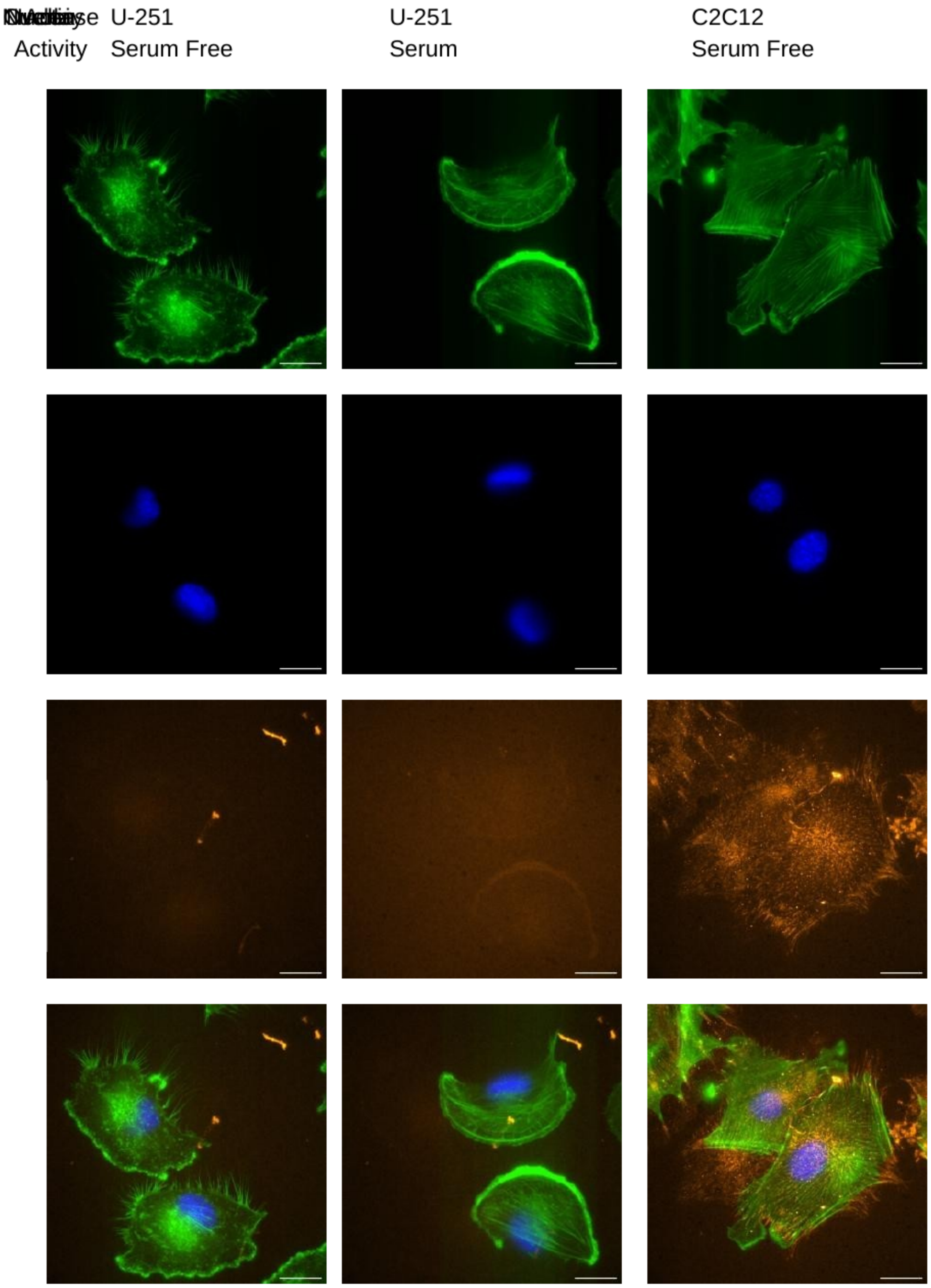


Supplementary Figure S10. Nuclease activity under representative RAD-TGT assay conditions. U-251 cells were plated on SNS-coated surfaces in serum-free or serum-containing medium, and C2C12 cells were plated in serum-free medium. Representative channels show actin, nuclei, SNS fluorescence, and overlays. Serum produced broadly distributed SNS signal around U-251 cells, whereas C2C12 cells generated strong localized SNS fluorescence beneath and around adherent cells even in serum-free medium, demonstrating contributions from both exogenous and cell-associated nucleases. Scale bars are as shown in the source images.

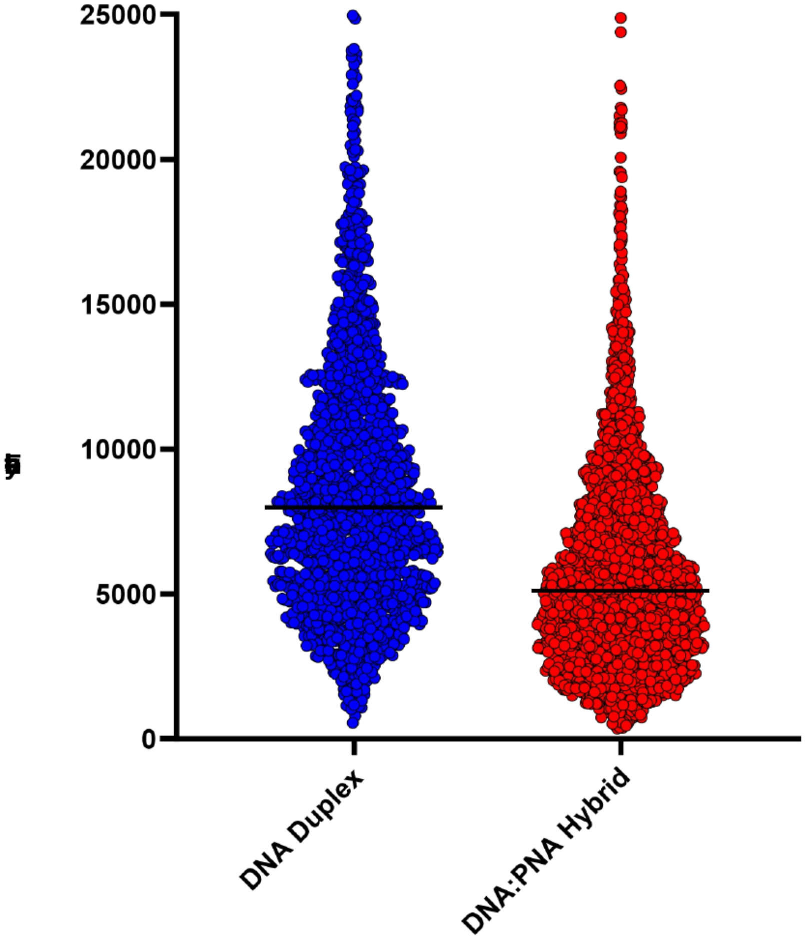


Supplementary Figure S11. RAD-TGT fluorescence obtained with DNA duplexes and DNA:PNA hybrid probes. Representative single-cell fluorescence distributions illustrate that DNA:PNA hybrid probes produce a lower absolute RAD-TGT signal than the corresponding DNA duplexes, consistent with the greater mechanical stability of the hybrid duplex. Horizontal lines indicate median fluorescence.

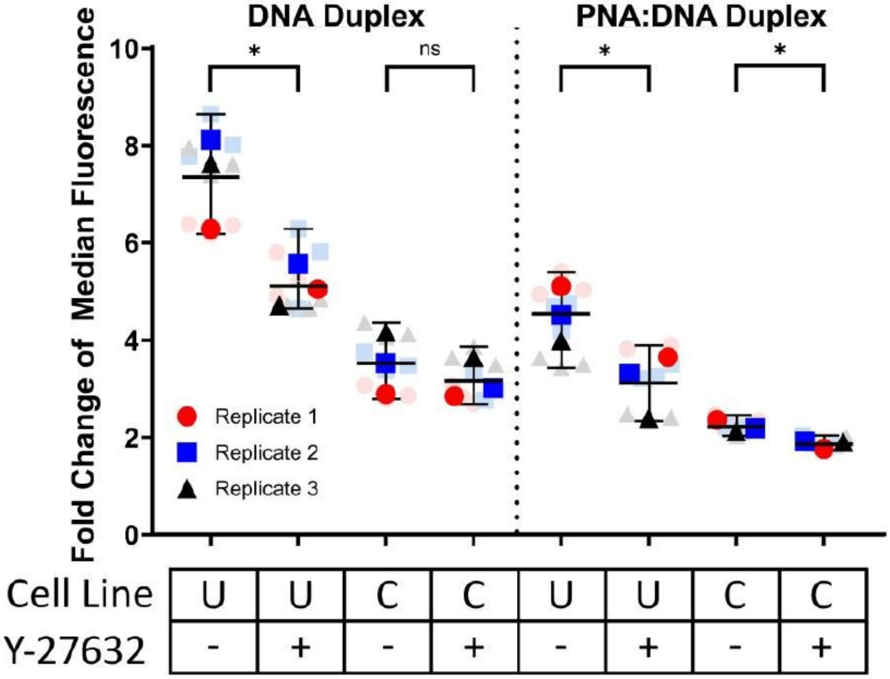


Supplementary Figure S12. DNA:PNA hybrids preserve detection of pharmacologically induced force changes in a high-nuclease environment. Low-nuclease U-251 (U) and high-nuclease C2C12 (C) cells were analyzed on shearing RAD-TGTs composed of DNA:DNA or DNA:PNA duplexes in the presence (+) or absence (-) of the ROCK inhibitor Y-27632. Data are shown as fold change in median fluorescence relative to the corresponding no-ligand control. Y-27632 reduced signal in U-251 cells with both probe types. In C2C12 cells, the drug-dependent reduction was not resolved with DNA duplexes but was recovered with DNA:PNA hybrids. Points represent three biological replicates; ns, not significant; *P < 0.05, unpaired Student t-test.

#### Supplementary Methods

##### Cell culture

Cell lines used for receptor-specific mechanophenotyping included U251, SKBR3, BT474, and SKOV3 cells. Cells were maintained under standard culture conditions, used at low passage number, and cultured under the medium and supplementation conditions appropriate for each line. Primary human immune-cell experiments were performed with human NK cells under the source, donor, isolation, and culture conditions used for the corresponding experiments. Primary human NK-cell isolation and culture. Peripheral blood products from healthy donors were obtained through Memorial Blood Bank (Minneapolis, MN). Peripheral blood mononuclear cells (PBMCs) were isolated by Ficoll-Paque density-gradient centrifugation, and NK cells were enriched from PBMCs by immunomagnetic negative selection using a human NK-cell isolation/enrichment kit (STEMCELL Technologies) according to the manufacturer's instructions. Studies involving human peripheral-blood products were reviewed and approved by the University of Minnesota Institutional Review Board under protocol 9709M00134. Donors provided informed consent according to Memorial Blood Bank policies, and blood products were deidentified prior to receipt by the laboratory.

##### Cloning of HUH-tagged recombinant antibodies

RVL4 (Addgene #104582) and RVL5 (Addgene #104583) antibody-expression plasmids were modified to encode the Wheat Dwarf Virus (WDV) HUH endonuclease tag followed by a C-terminal HA epitope. HUH-tag sequences were introduced at the C terminus of either the heavy- or light-chain constant region using restriction digestion and HiFi DNA assembly. Variable-region sequences for pertuzumab, which recognizes HER2, and K20, which recognizes β1 integrin, were obtained from SAbDab and/or the published literature and introduced into the corresponding heavy- and light-chain expression plasmids. Constructs used for the principal mechanophenotyping experiments encoded the WDV HUH tag on the antibody heavy chain.

##### Recombinant antibody expression and purification

Heavy- and light-chain plasmids were transiently co-transfected at a 1:1 plasmid ratio into Expi293F suspension cells using the ExpiFectamine 293 expression system. Conditioned medium was harvested 5–7 d after transfection and clarified by centrifugation. Recombinant antibodies were purified by Protein A affinity chromatography, eluted under acidic conditions with immediate neutralization, buffer exchanged, and further polished by size-exclusion chromatography as appropriate. Typical purified yields were approximately 10 mg antibody per liter of culture.

##### HUH-mediated oligonucleotide conjugation

HUH-mediated DNA conjugation was performed using single-stranded DNA substrates containing the WDV origin recognition sequence TAATATTAC. Unless otherwise indicated, HUH reactions were carried out in 50 mM HEPES, pH 8.0, 50 mM NaCl, and 1 mM $MnCl_2$ for 30 min at 37 °C. For analytical conjugation reactions, HUH-tagged antibody (3 µM) was incubated with DNA oligonucleotide (10 µM). Reaction products were analyzed by SDS-PAGE under reducing or nonreducing conditions as indicated. Under these conditions, the source experimental record reports >90% conjugation efficiency, with approximately 95% conjugation for both K20-HUH and pertuzumab-HUH fusions.

For preparation of antibody-functionalized tension probes, pre-annealed TGT duplexes were reacted with HUH-tagged antibody for 30 min at 37 °C. The source methods specify a 2:1 molar ratio of HUH-tagged antibody to TGT duplex.

##### SDS-PAGE analysis of HUH conjugation

HUH-mediated oligonucleotide conjugation and Protein G photocrosslinking products were resolved by SDS-PAGE using reducing or nonreducing sample conditions as appropriate to the experiment. Gels were used to assess mobility shifts associated with DNA conjugation and/or formation of covalent Protein G–IgG complexes.

#### Biolayer interferometry

Binding of recombinant HUH-tagged antibodies to their cognate antigens was measured using a BLItz biolayer interferometry instrument with anti-His biosensors. His-tagged recombinant human αVβ1 integrin or HER2 ectodomain was immobilized on the biosensor, and serial dilutions of antibody were analyzed using 120-s association and 120-s dissociation phases. Binding curves were analyzed in GraphPad Prism.

#### Production of photocrosslinkable Protein G–HUH adaptor

For adaptor-based antibody conjugation, the Protein G Fc-binding domain was engineered to contain an amber codon at Ala24 for incorporation of the photocrosslinkable unnatural amino acid 4-benzoyl-L-phenylalanine (BPA; also referred to as Bpa). The Protein G–HUH construct was co-expressed in BL21(DE3) Escherichia coli with the pEVOL-pBpF aminoacyl-tRNA synthetase/tRNA system. BPA was supplied during growth, and expression was induced with IPTG and arabinose. The resulting photocrosslinkable Protein G–HUH protein (clPG-HUH) was purified using the general purification workflow used for HUH fusion proteins.

#### Protein G photocrosslinking to Fc-containing proteins

clPG-HUH and Fc-containing antibody were combined at approximately a 1:1.5–2 molar ratio and incubated at room temperature to permit Fc binding. Samples were then illuminated on ice with 365-nm light to covalently trap the Protein G–Fc complex. A 20-min illumination time was selected for subsequent experiments because it provided a balance between efficient Fc photocrosslinking and retention of HUH-mediated DNA conjugation activity. Covalent complex formation and subsequent HUH activity were assessed by SDS-PAGE.

#### RAD-TGT probe assembly

Low- and high-threshold RAD-TGT probes were assembled using DNA duplex geometries corresponding to 12-pN and 54-pN rupture thresholds. Anchor and fluorescent ligand strands were mixed at a 1.1:1 molar ratio, with the fluorophore-containing strand limiting, in 10 mM Tris, pH 7.5, 50 mM NaCl, and 1 mM EDTA. Samples were heated to 98 °C for 5 min and cooled at room temperature for 1 h. The annealed duplexes were subsequently reacted with the indicated HUH-tagged ligand or antibody.

#### Preparation of antibody-functionalized RAD-TGTs

To generate receptor-specific RAD-TGTs, annealed TGT duplexes containing the WDV recognition sequence were covalently coupled to K20-HUH or pertuzumab-HUH through the sequence-directed HUH reaction. HUH-tagged antibody and TGT duplex were incubated for 30 min at 37 °C under the HUH reaction conditions described above. WDV HUH lacking a receptor-targeting ligand was used as a no-ligand control where indicated.

#### RAD-TGT surface functionalization

Glass-bottom 96-well plates were incubated with 80 µL per well of 100 µg/mL biotinylated BSA in PBS for 2 h at room temperature. Wells were washed twice with cold PBS, incubated with 100 µg/mL neutravidin for 30 min at room temperature, and washed again with PBS. Wells were then incubated with 80 µL of 1 µM prepared RAD-TGT overnight at 4 °C. Immediately before cell addition, wells were washed once with cold PBS and twice with Opti-MEM. For experiments requiring additional adhesion support, fibronectin was included in the biotinylated-BSA coating solution at 18.75 µg/mL.

#### RAD-TGT cell assay

Cells were plated onto functionalized RAD-TGT surfaces under the conditions indicated for each experiment and incubated at 37 °C and 5% $CO_2$. Receptor engagement and cell-generated force ruptured the tension-sensitive duplex and delivered the fluorescent oligonucleotide payload to cells. Following incubation, cells were recovered from the surface, washed as required, and prepared for flow-cytometric analysis.

#### RAD-TGT flow cytometry and feature extraction

Recovered cells were analyzed on a BD Accuri C6 Plus flow cytometer. Cells were gated using scatter, singlet, and viability criteria before analysis of the fluorescent RAD-TGT payload. Depending on the experiment, receptor-specific mechanical behavior was summarized using the percentage of RAD-TGT-positive cells, median fluorescence intensity (MFI), the high- to low-threshold signal ratio, and population heterogeneity measured by coefficient of variation (CV). For cancer-cell-line comparisons, measurements were normalized to the matched baseline or no-ligand control as specified for the corresponding analysis. High/low ratios were evaluated as candidate features but were not used as the primary multidimensional classifier because ratios become unstable when one component approaches background.

#### Multidimensional receptor-specific mechanophenotyping

Candidate mechanical features derived from β1-integrin- and HER2-targeting RAD-TGTs were compared across U251, SKBR3, BT474, and SKOV3 cells. The final two-dimensional representation used baseline-normalized median low-threshold β1-integrin engagement as the first coordinate and HER2 high-threshold signal heterogeneity (robust CV in the archived analysis output) as the second coordinate. Independent biological replicates were plotted individually or summarized as mean ± SEM, as indicated.

##### One-dimensional versus two-dimensional cell-line separation analysis

To quantify whether the combined receptor measurements improved discrimination among cell lines, all six pairwise comparisons among SKOV3, U251, SKBR3, and BT474 were evaluated. For each pair of cell lines, a standardized one-dimensional separation was calculated independently for the β1-integrin low-threshold median feature and the HER2 high-threshold CV feature. The better of these two constituent one-dimensional separations was designated the 'best 1D' separation for that cell-line pair. The two-dimensional separation was then calculated as the Euclidean combination of the two standardized coordinate separations:

$$\mathbf{d_2D = \sqrt{(d^2\_ITGB1 + d^2\_HER2)}}$$

This relationship is directly reproduced by the archived pairwise analysis output. For example, for SKOV3 versus U251, the standardized β1-integrin and HER2 separations were 1.575 and 3.097, respectively, giving a combined 2D separation of 3.474. The corresponding 2D values exceeded the best constituent 1D value for each of the six cell-line comparisons.

Across the six pairwise comparisons, the mean best-1D standardized separation was 6.177 and the mean combined 2D separation was 6.719, corresponding to a mean absolute gain of 0.541 standardized-separation units and a mean pairwise gain of 12.19%. A 20,000-iteration permutation analysis gave P = 0.00285 (reported as P = 0.0029). Bootstrap analysis gave a 95% confidence interval of 0.165–1.390 for the mean absolute gain and 1.31–16.32% for the mean pairwise percent gain.

> ***VERIFY BEFORE SUBMISSION:*** *The archived output files establish the pairwise distances, Euclidean 2D combination, 20,000 permutations, P value, and bootstrap confidence intervals. They do NOT preserve enough information to state with confidence (i) the exact denominator used to standardize each 1D mean difference or (ii) the exact label/sign-resampling algorithm used for the permutation and bootstrap tests. These two implementation details should be recovered from the original analysis script before submission rather than guessed.*

##### Pairwise separation values used in the 1D-versus-2D comparison

| Cell-line pair | β1 integrin 1D | HER2 CV 1D | Best 1D | Combined 2D |
|---|---|---|---|---|
| SKOV3 vs U251 | 1.575 | 3.097 | 3.097 | 3.474 |
| SKOV3 vs SKBR3 | 7.535 | 1.805 | 7.535 | 7.748 |
| SKOV3 vs BT474 | 5.395 | 0.803 | 5.395 | 5.455 |
| U251 vs SKBR3 | 4.839 | 3.198 | 4.839 | 5.800 |
| U251 vs BT474 | 3.955 | 4.259 | 4.259 | 5.812 |
| SKBR3 vs BT474 | 11.938 | 1.426 | 11.938 | 12.022 |

#### Multiplexed receptor-specific RAD-TGT measurements

For multiplexed measurements, HER2- and β1-integrin-targeting RAD-TGT probes were prepared with spectrally distinct fluorescent payloads and presented simultaneously. Following cell incubation and probe rupture, fluorescence in the two channels was measured by flow cytometry to generate a two-color receptor-specific mechanical signature for each cell. Mixed-cell or multi-cell-line samples were analyzed without requiring prior assignment of cell identity.

> ***VERIFY BEFORE SUBMISSION:*** *Insert the exact fluorophores, compensation procedure, probe concentrations/mixing ratio, and gating strategy used in the final multiplexed experiment.*

##### LFA-1 modulator experiments in primary human NK cells

Primary human NK cells were incubated with increasing concentrations of LFA-1-modulating antibodies before plating on ICAM-1-functionalized RAD-TGT surfaces. The antibody panel included the inhibitory antibody TS1/18, the activating antibody CBR LFA-1/2, and M24, which recognizes and stabilizes an activated/extended LFA-1 conformation. Following incubation on the RAD-TGT surface, cells were recovered and the delivered fluorescent payload was quantified by flow cytometry. Changes in RAD-TGT signal were plotted as a function of modulator concentration and fit to concentration-response curves. Concentrations producing half-maximal decreases or increases in the mechanical RAD-TGT response were designated mechanical $IC_{50}$ ($mIC_{50}$) or mechanical $EC_{50}$ ($mEC_{50}$), respectively, to distinguish these functional mechanical measurements from conventional binding-affinity or signaling-potency measurements.

##### Surface nuclease sensor assay

Surface nuclease sensors (SNS) were assembled from fluorophore- and quencher-containing DNA strands using the same general annealing workflow used for the tension probes. Probe-functionalized glass wells were prepared using biotinylated BSA and neutravidin. For SNS experiments, fibronectin was included in the biotinylated-BSA coating solution at 18.75 µg/mL to support cell adhesion. U251 cells were evaluated in serum-free and serum-containing medium to assess contributions from exogenous nuclease activity, whereas C2C12 cells were evaluated in serum-free medium to assess cell-associated nuclease activity. Cells were plated sparsely for microscopy and incubated at 37 °C and 5% $CO_2$. SNS fluorescence was imaged together with phalloidin-labeled actin and DAPI-labeled nuclei. CellProfiler was used to define cell-associated regions from the actin channel and to compare SNS fluorescence beneath cells with background regions.

##### DNA:PNA hybrid RAD-TGT assays

DNA:DNA duplexes were annealed in 10 mM Tris, pH 7.5, 50 mM NaCl, and 1 mM EDTA, whereas DNA:PNA hybrids were annealed in water. Anchor and ligand strands were combined at a 1.1:1 molar ratio with the fluorophore-containing strand limiting, heated to 98 °C for 5 min, and cooled at room temperature for 1 h. Shearing RAD-TGTs were functionalized with HUH-echistatin to engage RGD-binding integrins; HUH lacking a ligand fusion served as the matched negative control. U251 and C2C12 cells were treated with Y-27632 or matched DMSO vehicle for 30 min before plating. After 1.5 h on RAD-TGT surfaces, cells were collected and analyzed by flow cytometry. Results were quantified either as the percentage of cells exceeding the 99th percentile of the corresponding no-ligand control or as fold change in median fluorescence relative to the matched negative control. Biological replicates were compared using unpaired t-tests where indicated.

##### Statistical analysis

Biological replicates were treated as the unit of replication. Technical replicates were averaged within biological replicates where applicable. Data are reported as mean ± SEM unless otherwise indicated. Statistical tests were selected according to the experimental comparison and are specified in the corresponding figure legends. For DNA:PNA experiments, unpaired t-tests were used for the indicated two-group comparisons. Concentration-response experiments were analyzed by nonlinear regression in GraphPad Prism. The multidimensional mechanophenotyping analysis is described separately above.

#### Supplementary Table S1. Oligonucleotide and variable-region sequences

##### RAD-TGT oligonucleotides

| Oligonucleotide | Sequence / modification |
| --- | --- |
| 12-pN quencher anchor strand | 5′-/5IAbRQ/GGGTGGTCGCTGCGGGCC/3Bio/-3′ |
| 54-pN quencher anchor strand | 5′-/5IAbRQ/iBiodT/GGGTGGTCGCTGCGGGCC-3′ |
| Fluorescent ligand strand | 5′-GCTATAAACTCACCGTAATTTTTTGGCCCGCAGCGACCACCCTTT/3Cy5Sp/-3′ |
| WDV nonanucleotide recognition sequence | TAATATTAC |
| WDV ssDNA substrate reported in source table | TAATATTACCCCGCGTGG |

##### Antibody variable-region sequences

| Antibody | Chain | Amino-acid sequence |
| --- | --- | --- |
| Pertuzumab (HER2) | Light-chain variable region | DIQMTQSPSSLSASVGDRVTITCKASQDVSIGVAWYQQKPGKAPKLLIYSASYRYTGVPSRFSGSGSGTDFTLTISSLQPEDFATYYCQQYYIYPYTFGQGTKVEIK |
| Pertuzumab (HER2) | Heavy-chain variable region | EVQLVESGGGLVQPGGSLRLSCAASGFTFTDYTMDWVRQAPGKGLEWVADVNPNSGGSIYNQRFKGRFTLSVDRSKNTLYLQMNSLRAEDTAVYYCARNLGPSFYFDYWGQGTLVTVSS |
| K20 (β1 integrin) | Light-chain variable region | DIQLTQSPSSLSASLGGKVTITCKASQDINKYIAWYQHEPGKGPRLLIRYTSKLESGIPSRFSGSGSGRDYSFSISNLEPEDIATYYCLQYYNLWTFGGGTKLEIKRK |
| K20 (β1 integrin) | Heavy-chain variable region | QVQLQESGTELVRPGASVKLSCKASGYTFTDYYISWVKQRPGQGLEWIARIYPGSGNTFYNEKFKGKATLTAETSSNTAYMQLSSLTSEDSAVYFCAIYYGSGDYWGQGTTVTVSS |
